# Predicting antigen-specificity of single T-cells based on TCR CDR3 regions

**DOI:** 10.1101/734053

**Authors:** David S. Fischer, Yihan Wu, Benjamin Schubert, Fabian J. Theis

## Abstract

It has recently become possible to assay T-cell specificity with respect to large sets of antigens as well as T-cell receptor sequence in high-throughput single-cell experiments. We propose multiple sequence-data specific deep learning approaches to impute TCR to epitope specificity to reduce the complexity of new experiments. We found that models that treat antigens as categorical variables outperform those which model the TCR and epitope sequence jointly. Moreover, we show that variability in single-cell immune repertoire screens can be mitigated by modeling cell-specific covariates.

---

Antigen recognition is one of the key factors of T-cell-mediated immunity. The ability to accurately predict T-cell activation upon epitope recognition would have transformative effects on many research areas from in infectious disease, autoimmunity, vaccine design, and cancer immunology, but has been thwarted by lack of training data and adequate models. Although tremendous effort has been spent on elucidating the common rules that govern the TCR-pMHC interaction, it still remains elusive. The T-cell receptor (TCR) interacts with peptides immobilized on MHC multimers (pMHC) through its three complementarity determining region (CDR) loops of the α- and β-chain. The hypervariable loops CDR3α and CDR3β are most commonly aligned with the presented epitope^1^ and are hypothesized to be the main driver of T-cell specificity^2^. Due to lack of sufficient data, previous models for T-cell specificity were only based on the CDR3β loop^3,4,5^.

In this study, we exploit a newly developed single-cell technology that enables the simultaneous sequencing of the paired TCR α- and β-chain while determining the T-cell specificity to train multiple deep learning architectures modeling the TCR-pMHC interaction including both chains. The models include single-cell specific covariates accounting for the variability found in such data, thereby fully exploit the multiplicity of observations that can be easily sampled in single-cell screens. We show that models that include both α- and β-chain have a predictive advantage over models that only include the β-chain, while models fit on only a single chain still perform well. Interestingly, we further find that T-cell affinity imputation in a sample from a known donor is possible, enabling the assessment of the presence of disease-specific T-cells. Lastly, we anticipate a large number of single-cell studies involving T cells to exploit TCR-specificity as an additional phenotypic readout. To facilitate the usage of our predictive algorithms, we built the python package *TcellMatch* that hosts a pre-trained model zoo for analysts to impute pMHC-derived antigen specificities and allows transfer and re-training of models on new data sets.

## Results

### A joint deep learning model for alpha- and beta-chain, antigens, and covariates

Before the introduction of single-cell TCR reconstruction with coupled antigen binding detection via pMHCs (Fig. 1a), most paired observations of TCR and bound antigen only included the TCR β-chain, which are often found in entries of databases such as IEDB^6^ or VDJdb^7^. Here, we explore a data set based on single-cell pMHC capture in which paired α- and β-chain could be successfully reconstructed for 10,000s of cells and binding-specificity measured for 44 distinct pMHC complexes^8^. We designed a model to predict TCR-antigen binding based on α- and β-chain sequences and cell-specific covariates (Fig. 1b) using sequence-specific layer types such as recurrent layer stacks (bi-directional GRUs^9,10^ and bi-directional LSTMs^10,11^), stacks of convolutional layers^12^, self-attention^13^ layer stacks, and densely connected networks (Online Methods). We model binding events within a panel of antigens as a single- or multi-task prediction model through a vector of output nodes corresponding to antigens.

**Figure 1:**
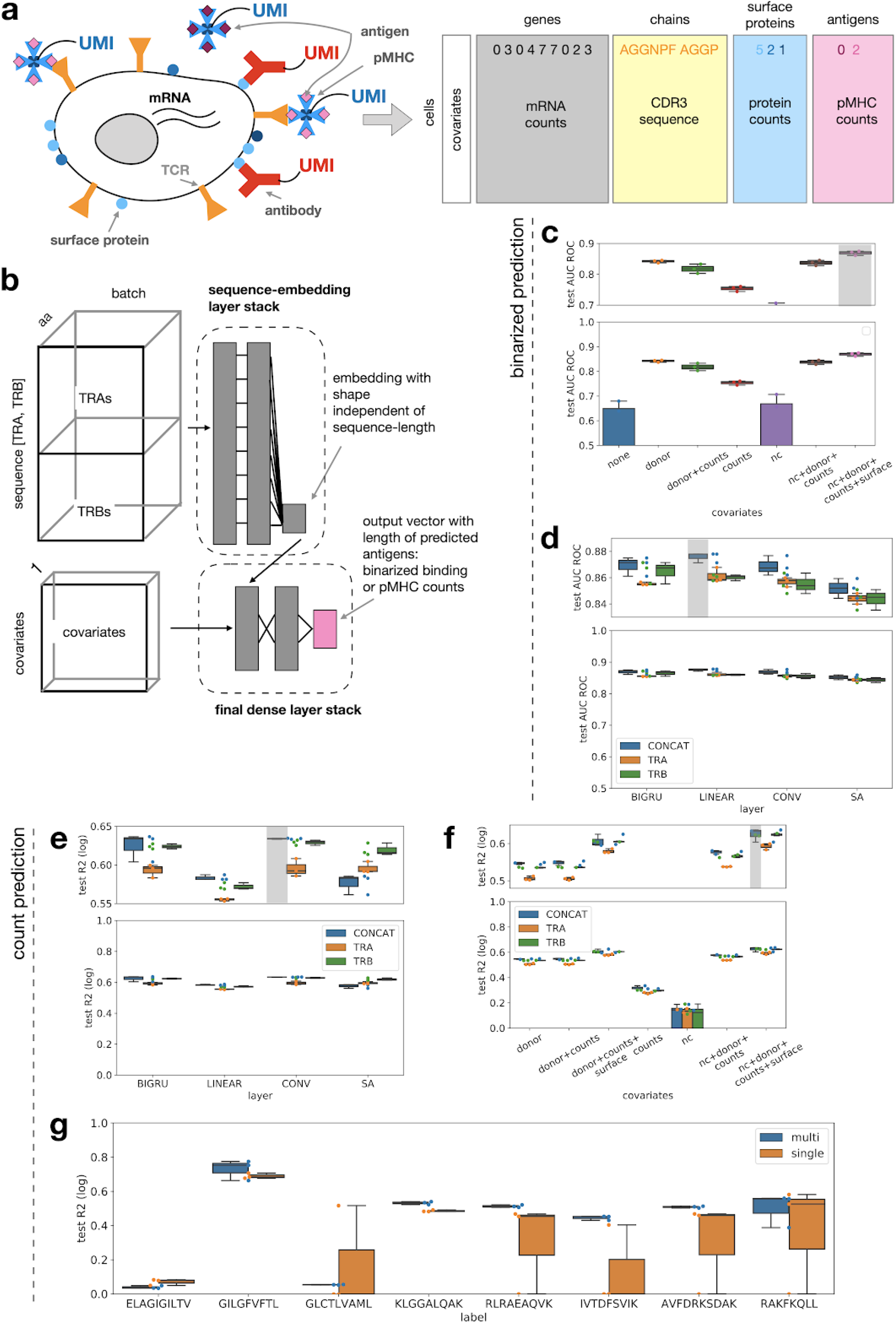
Deep learning models predict binding of TCRs to antigen panels. Grey boxes: Top performing model. Distributions shown as boxplots are across 3-fold cross-validation. (**a**) Concept of multimodal single-cell immune profiling experiment with RNA-seq, surface protein quantification, bound pMHC quantification, and TCR reconstruction. (**b**) Categorical TcellMatch model: A feed-forward neural network to predict a vector of antigen specificities of a T-cell based on the CDR3 TCR α- and TCR β-chain sequences. Grey boxes: layers of the feed-forward network. (**c**) Covariates improve sequence-based binding accuracy prediction. AUC ROC test: Area-under the receiver operator characteristic curve on the test set for the binary binding event prediction task. The top panel is a zoom into an informative region of the y-axis. *counts*: total mRNA counts, *nc*: negative control pMHC counts, *surface*: surface protein counts. (**d**) Antigen binding prediction based on TCR CDR3 sequences is improved by modeling α- and β-chain. *BIGRU*: bi-directional GRU model, *SA*: self-attention model, *CONV*: convolution model, *LINEAR*: linear model. (**e**) Sequence-encoding layer types out-perform linear models on pMHC count prediction if donor and size factors are given as covariates. *BIGRU*: bi-directional GRU model, SA: self-attention model, *CONV*: convolution model, *LINEAR*: linear model. (**f**) Performance of bi-directional GRU models that predict pMHC counts directly is best if covariates and both TCR chain are modeled. *test MLSE2*: mean logarithmic squared error on the test set, *test R2 (log)*: test R2 on log-transformed test data. (**g**) Multitask models outperform separate single-task model on pMHC count prediction by antigen. *multi*: multitask model, *single*: single-task model. All boxplots: The center of each boxplots is the sample median, the whiskers extend from the upper (lower) hinge to the largest (smallest) data point no further than 1.5 times the interquartile range from the upper (lower) hinge.

### Cell-specific covariates improve binding event prediction

Single-cell T-cell affinity screens feature multiple effects that confound the binding observation. Firstly, one would expect the donor identity to affect the TCR structure if donors vary in their HLA genotype. We compared models with and without a one-hot encoded donor identity covariate to establish the impact of these donor-to-donor differences. Firstly, we removed putative doublets from the data set (Online Methods, Supp. Fig. 1). To remove effects from strong class imbalance, we only considered the 8 antigens in the pMHC CD8+ T-cell data set that had at least 100 unique, non-doublet clonotype observations (Supp. Fig. 2a,b). The total data set size was 91,495 unique, non-doublet observations (cells) across four donors. We found that the performance of models without donor information varies strongly and is much worse than the performance of models with donor covariates (Fig. 1c). The initial amino acid embedding did not have a strong effect on the results (Supp. Fig. 3). These categorical models also performed well on data derived from the public databases (IEDB^6,7^ and VDJdb^7^) even though there were no corresponding covariates present (Supp. Fig. 4).

**Figure 2:**
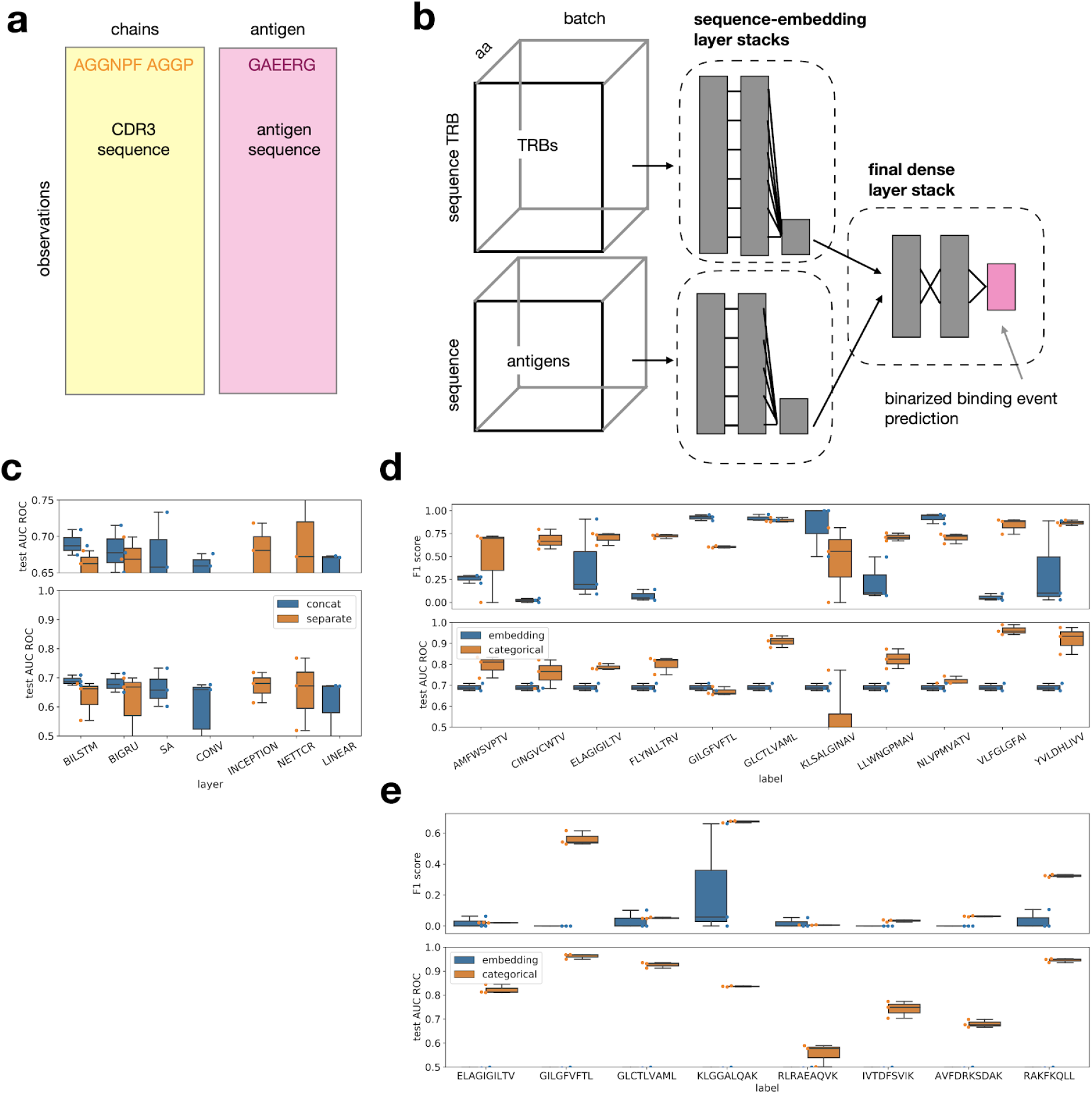
Deep learning models predict affinity of TCRs to sequence-encoded antigens. Distributions shown as boxplots are across 3-fold cross-validation. (**a**) The databases IEDB and VDJdb contain pairs of TCRs and antigens that were found to be specific to each other and are curated from many different studies. Supervised model that predict binding events can be trained on such data but also require the assembly of a set of negative observations (Online Methods). (**b**) Antigen-embedding TcellMatch model: A feed-forward neural network to predict a binding event based on TCR CDR3 sequences and antigen peptide sequence. Grey boxes: layers of the feed-forward network. (**c**) Different sequence encoding layer types perform similarly well on binding prediction based on TRB-CDR3 and antigen sequence. *CONCAT*: Models in which TRB CDR3 sequence and antigen sequence are concatenated, *SEPARATE*: Models in which TRB CDR3 sequence and antigen sequence are embedded by a separate sequence encoding layer stacks. *BILSTM*: bi-directional LSTM model, *BIGRU*: bi-directional GRU model, SA: self-attention model, *CONV*: convolution model, *INCEPTION*: inception-type model, *NETTCR*: NetTCR model^3^, *LINEAR*: linear model. (**d, e**) Antigen-wise categorical models outperform models that are built to generalize across antigens on high-frequency antigens in IEDB (**d**) and on overlapping antigens between IEBD and 10x CD8^+^ data (**e**). *embedding*: models that are embedding the antigen sequence and can be run on any antigen (Fig. 2a), *categorical*: Antigen-wise categorical models that do not have the antigen sequence as a feature (Fig. 1b). All boxplots: The center of each boxplots is the sample median, the whiskers extend from the upper (lower) hinge to the largest (smallest) data point no further than 1.5 times the interquartile range from the upper (lower) hinge.

The identification of binding events based on single-cell RNA-seq libraries is liable to false negatives due to low capture rate of RNAs. In standard single-cell RNA-seq processing, such effects are often rectified through normalization. We investigated, whether such normalization factors and negative control pMHC counts are useful predictors of a false negative binding event: We compared models only considering the donor identity covariate and models that also included a scaled total mRNA count covariate and ones that contained negative control count covariates (Online Methods). Across all architectures, models that accounted for the total mRNA count or the negative control counts of a cell performed better than models that did not do so, suggesting that false-negative correction is feasible (Fig 1c). We could also identify a predictive advantage of models that accounted for the cell type encoded by surface protein counts (Fig. 1c). We hypothesize that the surface protein counts can be used to embed cells based on their membrane surface structure which in turn could correlate with the number of TCRs on the cell surface. Accordingly, the integration of surface proteins in the model could correct for variance induced by cell-specific TCR availability. The overall top-performing model accounted for donor, total counts, negative control counts and surface protein counts (Fig. 1c).

### Co-modeling alpha- and beta-chain improves binding event prediction

We compared prediction performance between models fit using one TCR CDR3 chain (“TRA-only”, or “TRB-only”), to models fit to the concatenated TRB and TRA chains (“TRA+TRB”) to evaluate the additional information that one can gain by using both the TRA and TRB chain. We found that TRA+TRB models were consistently better than TRA-only and TRB-only models across most layer types if basic single-cell covariates were included in the prediction (Fig. 1d). We found that self-attention, recurrent and convolutional neural networks performed similarly to linear models (Fig. 1d). This suggests that antigen-specificity of a α- and β-chain pair can be well represented as a sequence motif problem in which the sequence motif has a fixed position on the CDR3 sequence.

### Continuous binding affinities can be predicted based on pMHC counts

In single-cell-based studies, antigen-binding events are measured based on the number of bound pMHCs of the target antigen and bound negative control antigens (Fig. 1a). The raw data describing the binding event is not a binary signal but lies in the positive integer space (count data). This opens up the possibility to not only model binding events (binarized signal) but also binding affinity, which enables the prioritization of highly affine epitopes for vaccination and the rational design of TCR sequences binding a specific antigen. We fit models that were similar in structure to the models dedicated to binarized binding event prediction on covariates and TCR CDR3 sequences to predict pMHC counts per cell (Fig. 1b). Again, TRA+TRB models outperformed TRA-only and TRB-only models across layer types (Fig. 1e). Covariates improved predictive power and models with donor, total counts, negative control pMHC and surface proteins count covariates performed best again (Fig. 1f).

Low-affinity binding events that are not captured in the discretized binding data but may be represented in the pMHC counts. Such low-affinity events may contain information about antigen-antigen similarities and therefore about output-space correlations, which can be exploited by multi-task supervised learning. Indeed, we found that multi-task models outperformed single-task models on six out of eight antigens modelled (Fig. 1g). An alternative interpretation of the improved performance of multi-task models is their ability to learn better de-noised low-dimensional representations of TCR sequences, through the integration of more diverse training data.

### Models with sequence-space embedding of antigens are outperformed by categorical models

Binding events in the databases such as IEBD^6^ or VDJdb^7^ (Fig. 2a) have previously been modeled based on a learned embedding of the antigen amino acid sequence^3^ (Fig. 2b). Here, we investigate whether such antigen-embedding models outperform simple, antigen-wise logistic models of binding events and whether they can generalize to unseen antigens.

Firstly, we benchmarked models with different layer types that predict a binding event based on sequence embeddings of the antigen and TCR β-chain. Previously, a specific single-layer motif-based architecture was proposed for this task^3^. We found that all common sequence-embedding layer types, are able to perform this prediction and that recurrent neural networks perform best in terms of model uncertainty (Fig. 2c).

In contrast to the categorical approach before, generalization across antigen sequences cannot easily be performed based on sequence motif recognition. We hypothesized that antigen-embedding models could learn a matching of seen antigens to TCRs within which the prediction problem can be broken down to a TCR motif-detection problem. In this setting, antigen-wise models that identify the antigen categorically in the output should be superior as they do not have to solve the matching problem. We found that antigen-wise categorical models have a better predictive performance on the antigens they were trained on than sequence-embedding models, on both the IEDB and pMHC CD8+ T-cell data set (Fig. 3d,e). We conclude that the previously proposed antigen sequence-embedding models are currently suboptimal for binding prediction on seen antigens.

**Figure 3:**
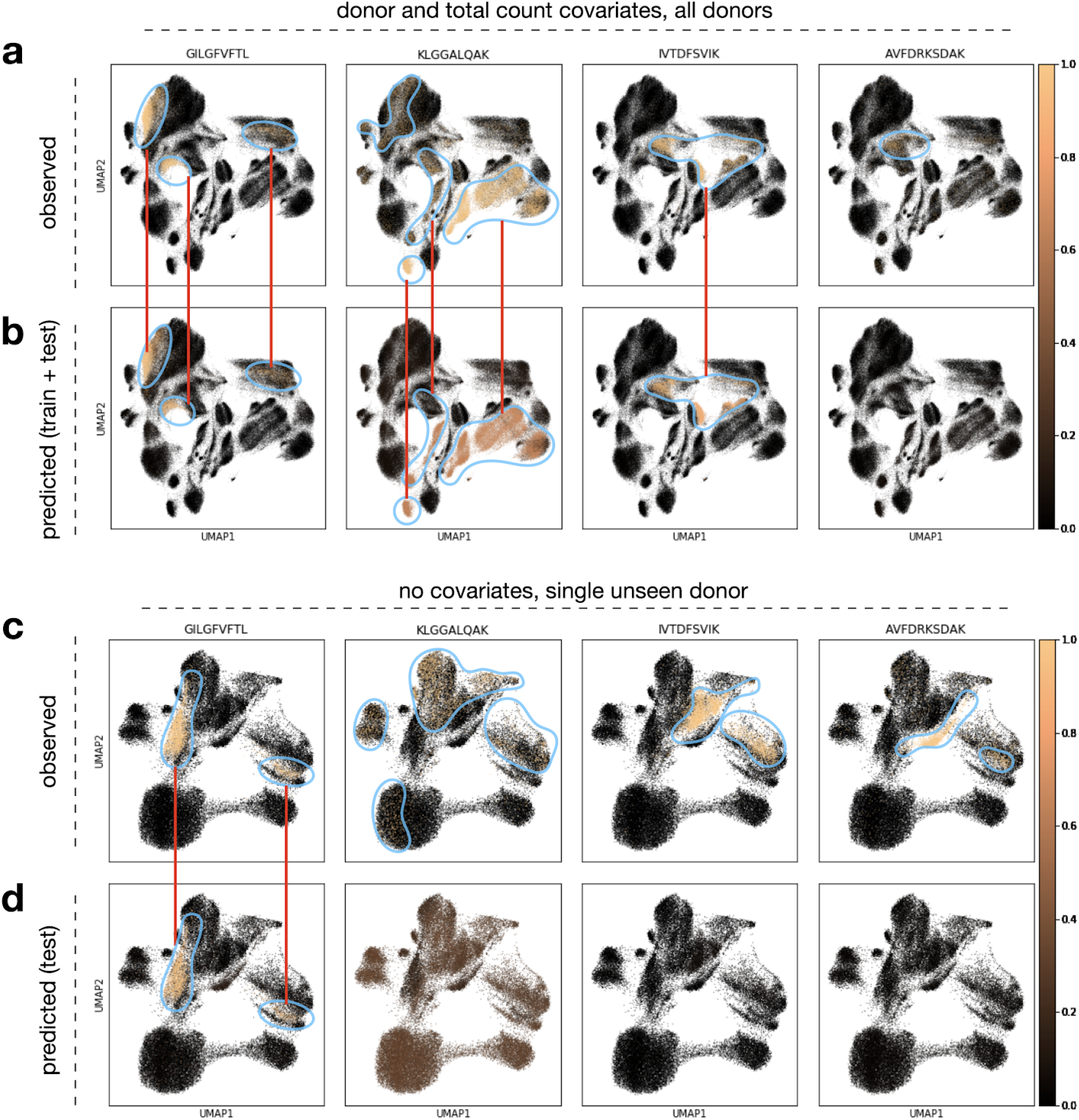
Imputed antigen specificity labels enrich single-cell RNA-seq workflows on T cells by an additional phenotype. (**a-d**) UMAP with observed (**a, c**) and predicted (**b, d**) labels. (**a, b**) The cells in the UMAP are the cells from all donors (train and validation data, n=189,512), the model was fit with donor and size factor covariates. (**c, d**) The cells in the UMAP are the cells from a validation donor (n=46,526), the model was fit without covariates.

Given that current datasets do not adequately cover the antigen space, we tested the current potential of sequence-embedding models to generalize to unseen antigens. This task cannot be covered by models that treat antigens as categories. Firstly, we trained models on a subset of high-frequency antigens from IEDB and tested on low-frequency antigens from IEDB and found the IEDB trained models do not generalize well to these antigens (Supp. Fig. 5a). Secondly, we used a subset of observations of VDJdb with antigens not overlapping to IEDB as a test set (Supp. Fig. 5b) and found that models trained on antigens occurring in IEDB do not generalize well to these antigens either. Thirdly, models trained on IEDB performed poorly on predicting binding in the pMHC CD8^+^ T-cell data (Supp. Fig. 5c). Thus, we cannot find evidence in the current TCR databases that extrapolation in the antigen space is possible based on current numbers of sampled antigens.

### Imputation of antigen-specificity of T-cells adds phenotypic information to single-cell studies

We showed that antigen specificity can be predicted based on TCR sequences from single-cell data. The training of such models requires single-cell experiments with pMHC binding detection. The inclusion of pMHC binding detection in an experiment increases the sequencing and reagent costs compared to CDR3 sequencing only experiments; this will be especially drastic in assays with many different antigens. Therefore, we believe that imputation of antigen specificity based on pre-trained models will be a valuable alternative to including pMHCs in T-cell assays. All models discussed above can be used for the purpose of imputation. We found that antigen specificity imputation can give interpretable results in T-cell subpopulations identified based on the transcriptome (Fig. 3). The observed labels are enriched in sub-regions of the transcriptome space (Fig. 3a,c) which can be recovered in multiple cases based on the predicted labels (Fig. 3b,d).

## Discussion

Our results demonstrate the benefit of jointly modeling the TCR α- and β-chain while accounting for single-cell variability through cell- and donor-specific covariates for T-cell specificity prediction. Most importantly, we found models that treat antigens as categorical outcome variables outperform those that model the TCR and antigen sequence jointly. Our results suggest that T-cell specificity can be predicted in an HLA genotype-specific fashion and thereby pave the way for research and development on all HLA types, beyond the commonly investigated type HLA-A*02:01. Generalization to unseen antigens with sequence-embedding models is currently challenging, but will become an important future research topic once screens with larger pMHC panels become available. Lastly, we showed that pMHC counts can be modeled as a measure of continuous binding affinity and that multi-task models outperform single-task models in this setting, paving the way for the integration of large pMHC panels in single models.

T-cell specificity complements standard immunological single-cell RNA-seq studies, and can be used to uncover subpopulations that are expected to be activated during disease or used as an indicator of antigen presence in a tissue. Consequently, we believe that the computational imputation of T-cell specificity will become an important tool for immunologically focused single-cell RNA-seq experiments. Imputation will reduce experimental complexity and costs and will also offer unbiased specificity metrics that are not liable to errors in the pMHC panel choice. Such prediction models can also be directly applied to immunophenotyping by screening for TCRs that interact with known viral or cancer neoepitopes, enabling the characterization of a patient’s immunological state and the stratification of subpopulations that are amenable for antigen-specific immunotherapies. Continuous T-cell binding affinity models would enable the possibility of rational in silico TCR design, accelerating the development of TCR-based biologics.

**Supp. Fig. 1:**
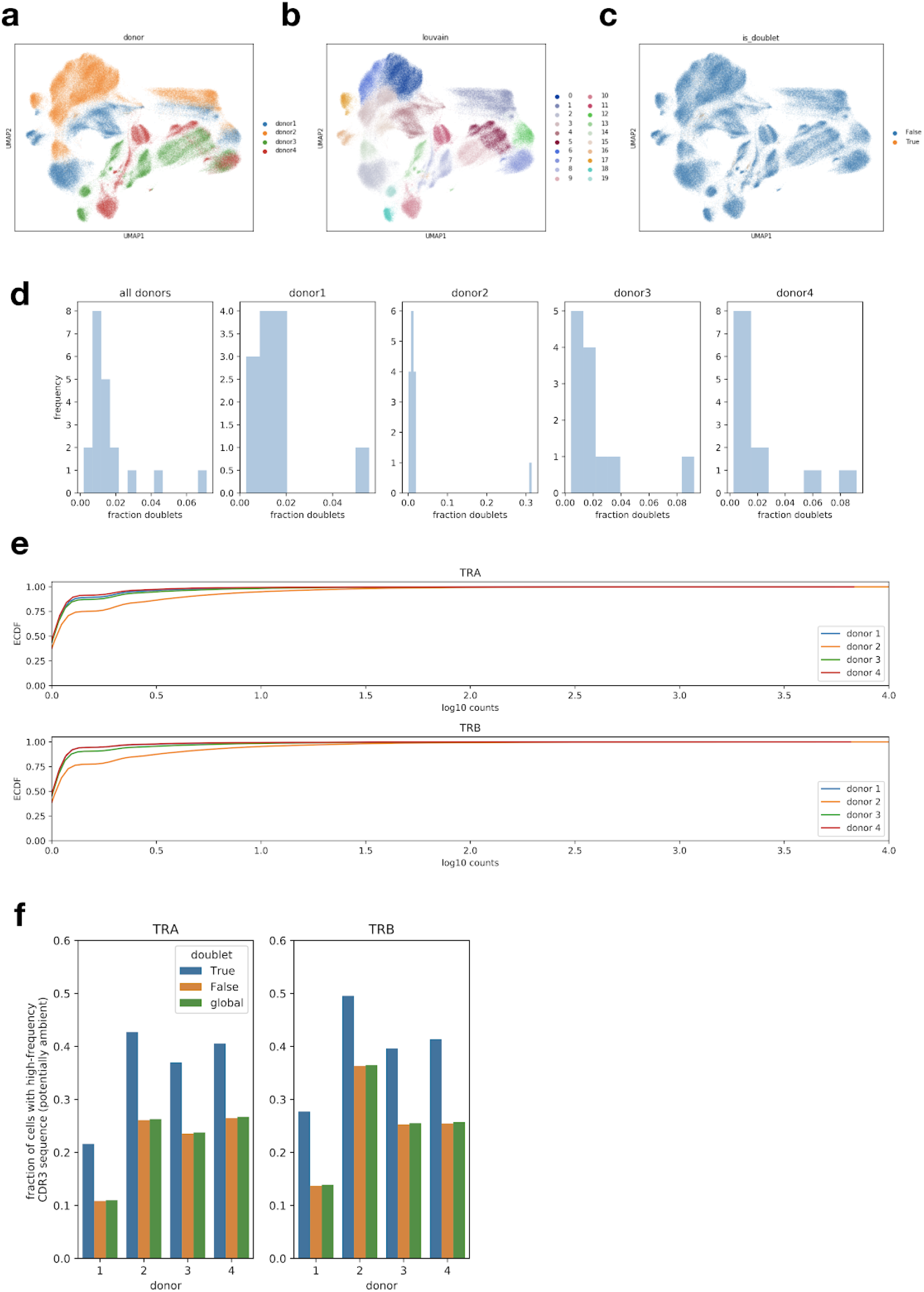
Cellular doublet identification based on non-unique TCR chain reconstructions. (**a-c**) UMAP of CD8^+^ T cells from all donors (n=189,512) computed based on the transcriptome with (**a**) donor identity, (**b**) Louvain cluster and (**c**) inferred doublet state superimposed. (**d**) Distribution of fractions of doublet out of all cells per clustering computed for each donor and for all clustering computed across all donors. (e) Empirical cumulative density function (ECDF) of the number of T cells that have a given CDR3 TCR sequence by chain and donor. *log10 counts* on the x-axis are the base 10 logarithm of the number of T cells for a given CDR3 sequence. (**f**) The fraction of cells that contain high-frequency CDR3 sequences which occur in more than 50 clonotypes. These high-frequency sequences are defined separately for each donor and may partially represent sequences derived from ambient molecules (Online Methods). *True*: is doublet, *False*: is not doublet, *global*: All cells, doublets, and non-doublets.

**Supp. Figure 2:**
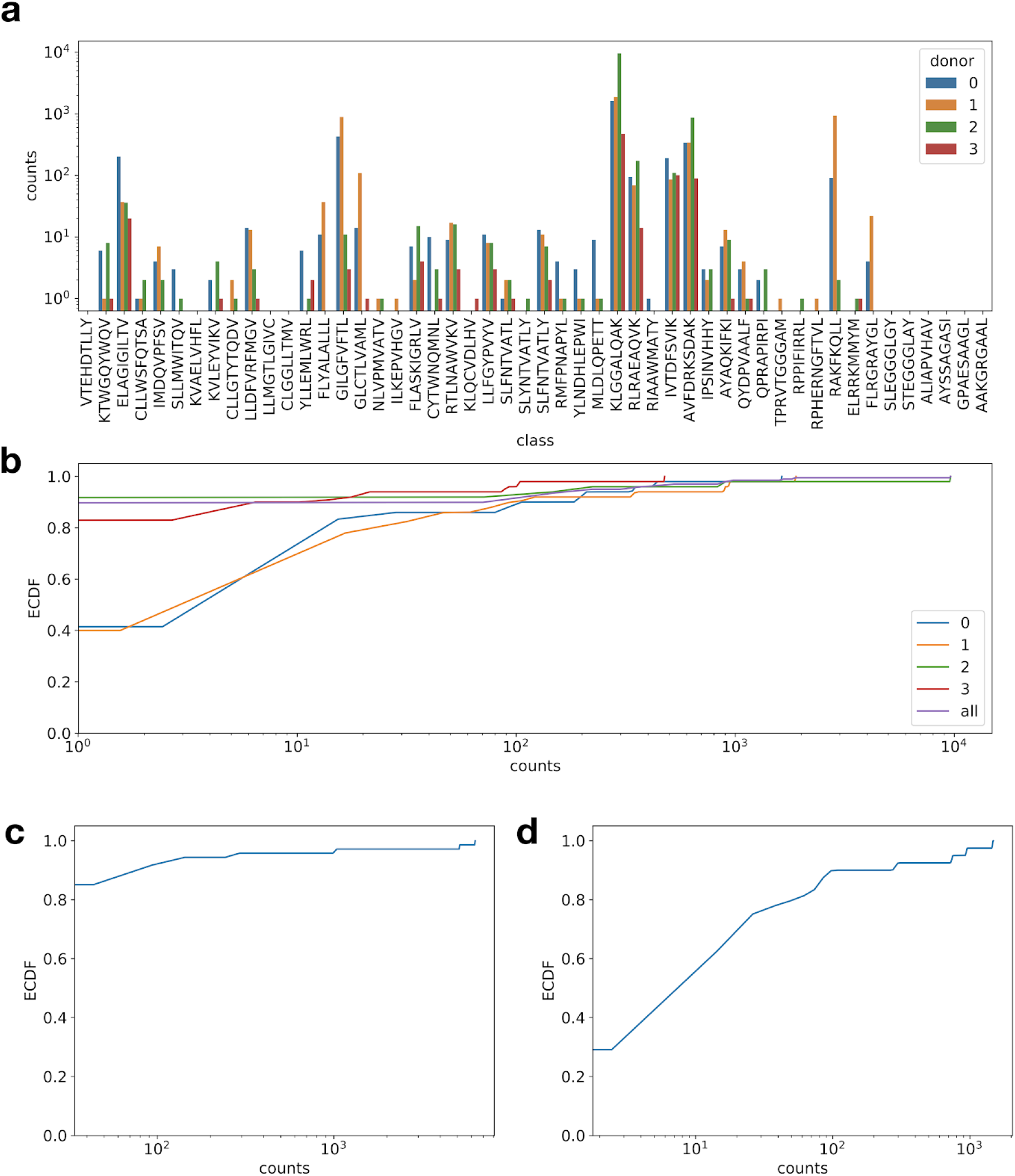
Number of unique TCR observations per antigen. (**a**) Histogram with the number of TCR clonotypes by antigen and donor for 10× CD8^+^ T-cell immune repertoire data, (**b-d**) Empirical cumulative density function (ECDF) of number of clonotypes (counts) per antigen for 10× CD8^+^ T-cell immune repertoire data (**b**), IEDB (**c**) and VDJdb (**d**).

**Supp. Figure 3:**
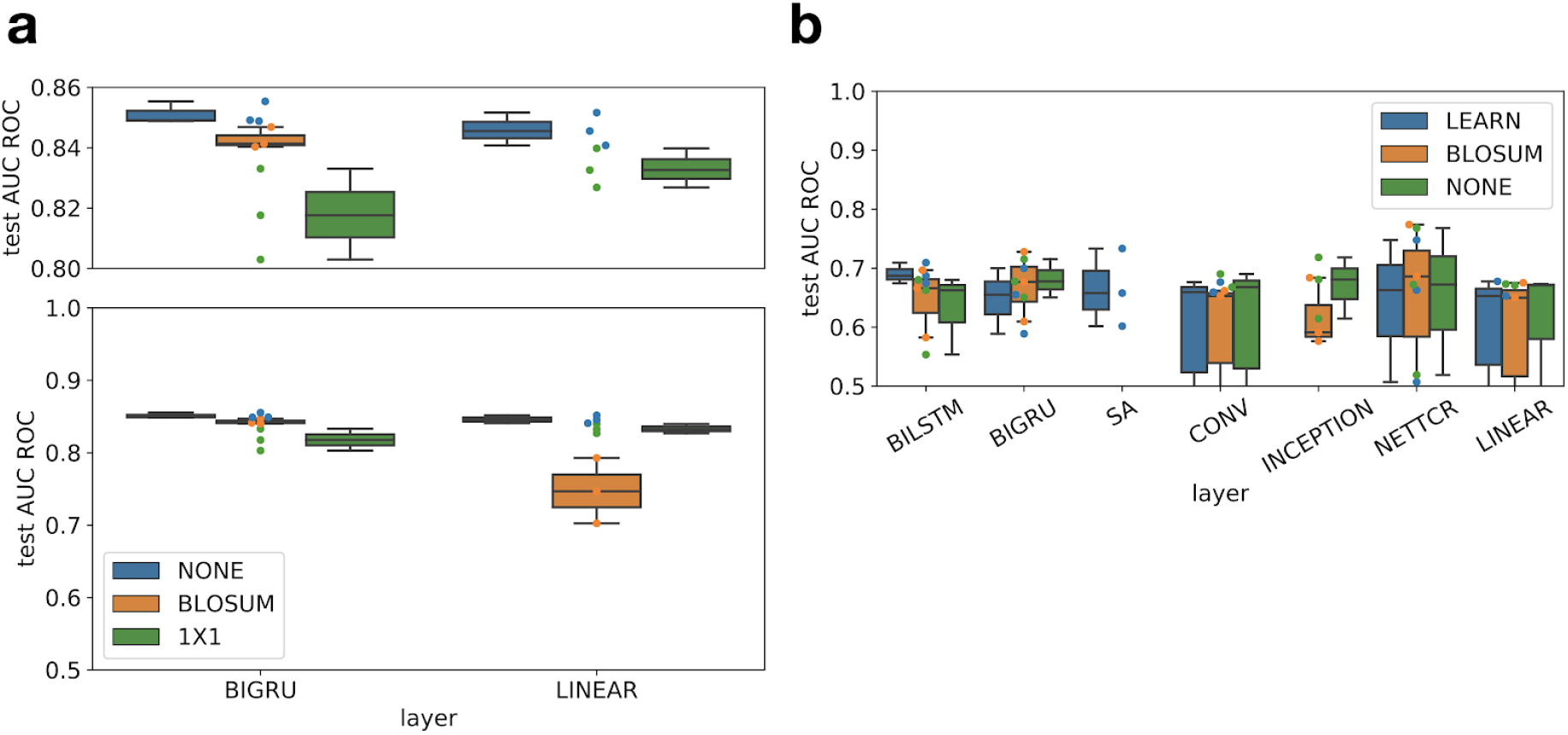
Amino acid embedding choice does not strongly affect model performance. Distributions shown as boxplots are across 3-fold cross-validation. (**a, b**) Comparison of model performance given multiple initial amino acid embeddings for models with antigen identity encoded in the output (**a**) and for models with sequence embedding of the antigen in the feature space (b). *BLOSUM*: BLOSUM52 embedding, *NONE*: one-hot encoding, *1×1* 5-dimensional 1×1 convolution on top of BLOSUM52 embedding that is learned at training time. All boxplots: The center of each boxplots is the sample median, the whiskers extend from the upper (lower) hinge to the largest (smallest) data point no further than 1.5 times the interquartile range from the upper (lower) hinge.

**Supp. Figure 4:**
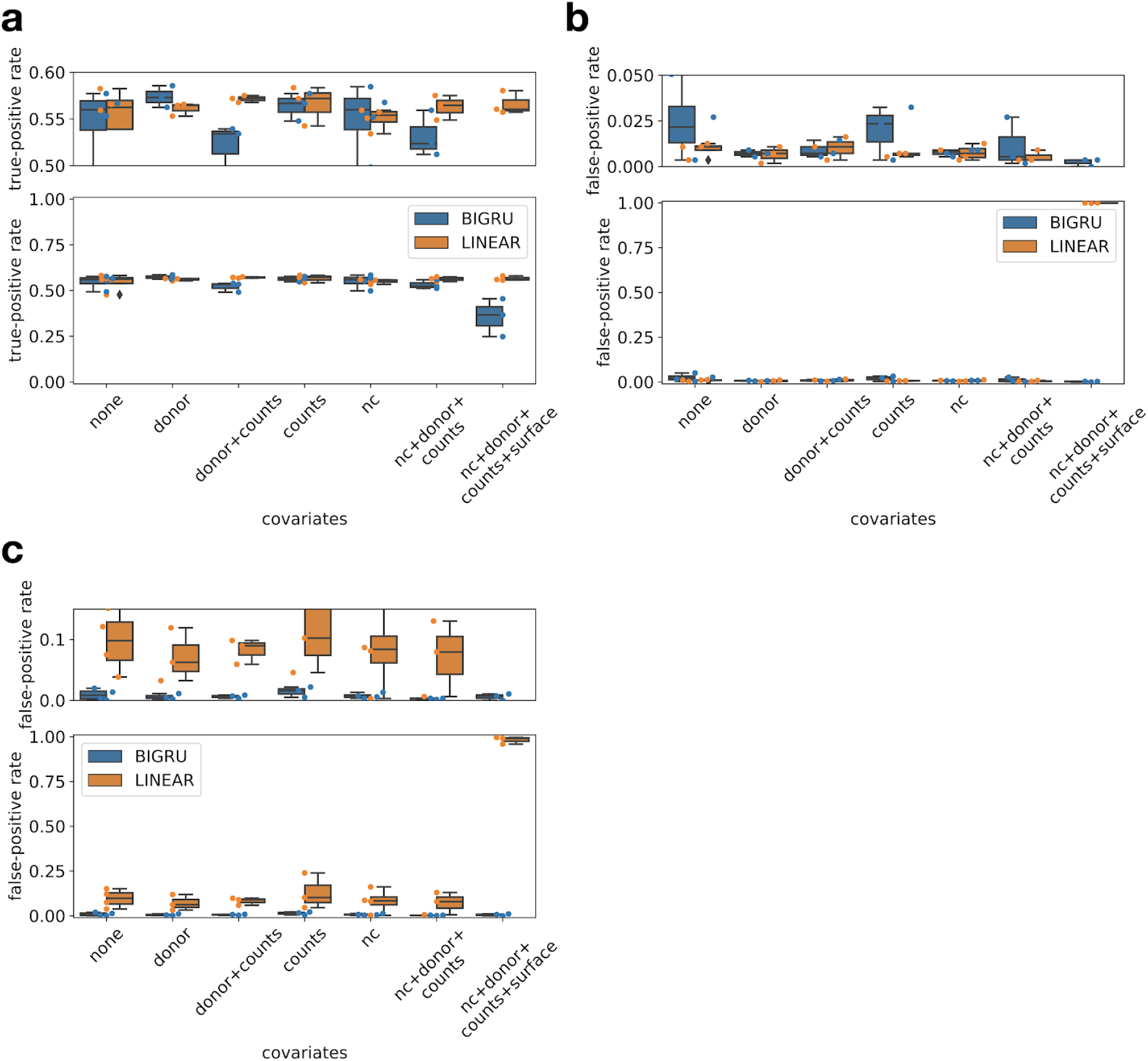
Validation of categorical models learned on pMHC CD8+ T-cell data on IEDB and VDJdb. Distributions shown as boxplots are across 3-fold cross-validation. (**a**) True-positive rate of best performing model by layer type and covariate setting on VDJdb entries with antigens that occur in the pMHC panel. All observations in this set should be predicted as positive for one of the categories of the model. *counts*: total mRNA counts, *nc*: negative control pMHC counts, *surface*: surface protein counts. (**b, c**) The false-positive rate of best performing model by layer type and covariate setting on VDJdb (**b**) and IEDB (**c**) entries with antigens that do not occur in the pMHC panel. All observations in this set should be predicted as negative (not binding any antigen of the panel). All boxplots: The center of each boxplots is the sample median, the whiskers extend from the upper (lower) hinge to the largest (smallest) data point no further than 1.5 times the interquartile range from the upper (lower) hinge.

**Supp. Figure 5:**
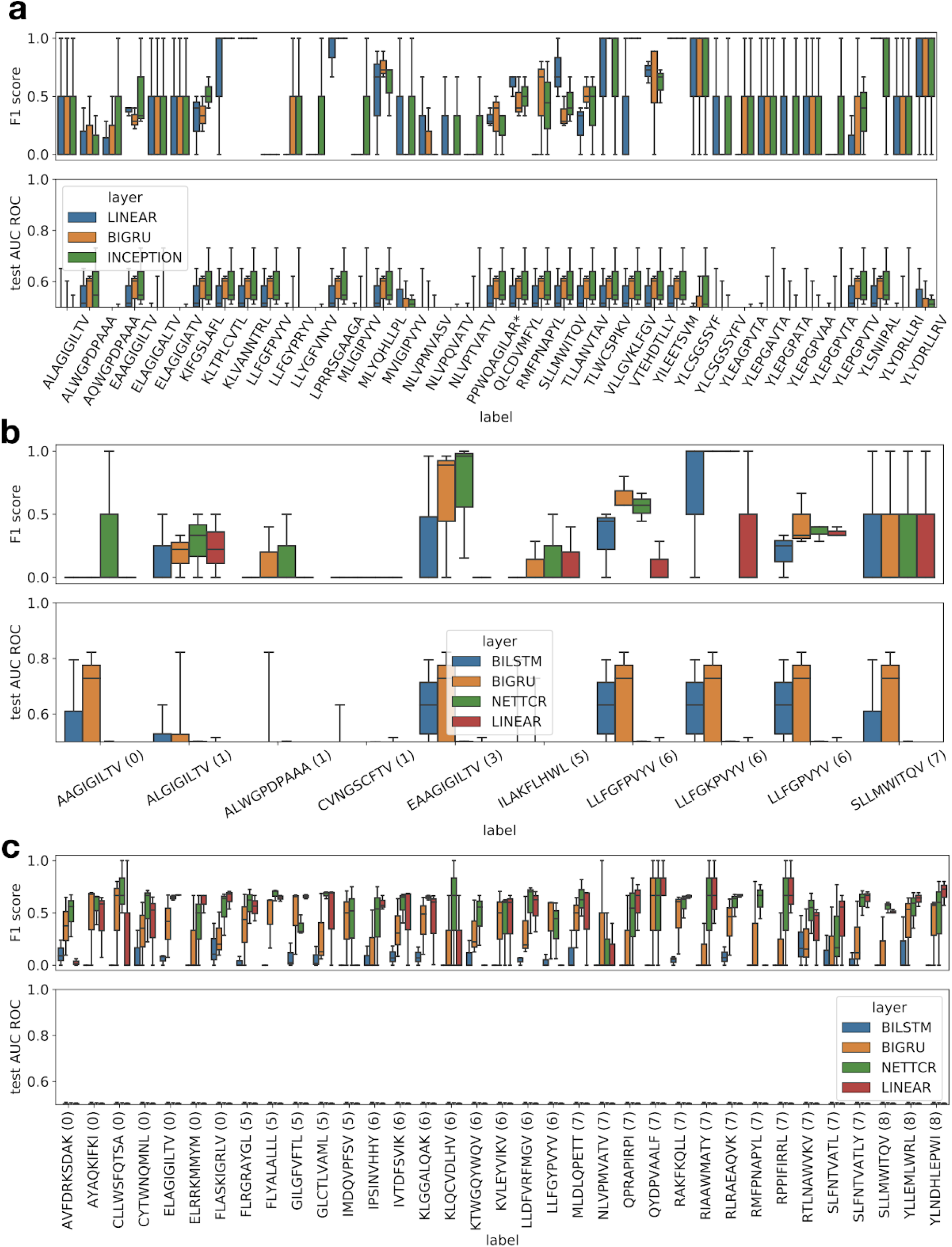
Models that embed antigen sequences to predict binding events cannot generalize well to unseen antigens. *BIGRU*: Models trained with bidirectional GRUs as sequence-embedding layers. *NETTCR*: NetTCR-like model. *LINEAR*: Models trained with a single densely connected layer as a sequence-embedding layer. *test AUC ROC*: Area-under the receiver operator characteristic curve on the test set for the binary binding event prediction task, *F1 score*: F1 score on binary predictions on the test set. Distributions shown as boxplots are across 3-fold cross-validation. (**a**) Models trained on antigens in IEDB cannot generalize to unseen low-frequency antigens in IEDB. (**b**) Models trained on all antigens from IEDB data cannot generalize to unseen antigens in VDJdb. (**c**) Models trained on all antigens from IEDB data cannot generalize to unseen antigens in 10x CD8+ data set. All boxplots: The center of each boxplots is the sample median, the whiskers extend from the upper (lower) hinge to the largest (smallest) data point no further than 1.5 times the interquartile range from the upper (lower) hinge.

## ACKNOWLEDGEMENTS

None.

## Data and code availability

The Python package *TcellMatch* will be available from GitHub (https://github.com/theislab/tcellmatch). All data is publicly available and was downloaded and processed as described in the Online Methods.

## Online Methods

### Feed-forward network architectures

Here, we describe proposed architectures of the models that predict antigen specificity of a T-cell receptor (TCR) based on the CDR3 loop of both α- and β-chain and on cell-specific covariates. Note that specificity determining influences of CDR1 and CDR2 loops^14–16^ and distal regions^17,18^ have been demonstrated as well but were not measured in the single-cell pMHC assay. All networks presented contain an initial amino acid embedding, a sequence data embedding block and final densely connected layer block.

#### Amino acid embedding

The choice of initial amino acid embedding may impact data and parameter efficiency of the model and therefore may impact predictive power of models trained on data sets that are currently available. We used one-hot encoded amino acid embeddings, evolutionary substitution-inspired embeddings (BLOSUM) and learned embeddings. The learned embeddings were a 1×1 convolution on top of a BLOSUM encoding and were prepended to the sequence model layer stack. Here, channels are the initial amino acid embeddings (we chose BLOSUM50) and filters are the learned amino acid embedding. This learned embedding can reduce the parameter size of the sequence model layer stack. All fits presented in the manuscript other than in Supp. Fig. 3 are based on such a learned embedding with 5 filters. We anticipate sequence-based embeddings to gain relevance in the context of extrapolation across antigens in the future. Here, parameter efficiency in the sequence models will play an important role and the 1×1 convolution presented here is an intuitive first step into this direction.

#### Sequence data embedding

We screened multiple layer types in the sequence data embedding block: Recurrent layers (bi-directional GRU and LSTM), self-attention, convolutional layers (simple convolutions and Inception-like), and densely connected layers as a reference. Recurrent layer types and self-attention layers have been previously useful for modeling language^13^ and epitope^19^ data. Convolutional layer types have been useful for modeling epitope^20,21^ and image^12^ data. The sequence-model layers retain positional information in subsequent layers and can thereby build an increasingly abstract representation of the sequence. To achieve this on recurrent networks, we chose the output of a layer to be a position-wise network state which results in an output tensor of size (batch, positions × 2, output dimension) for a bi-directional network. This position-wise encoding occurs naturally in self-attention and convolutional networks. We did not use feature transforms with positional signals^13^ on the self-attention networks, so that the network has no knowledge of the original sequence-structure but can still retain inferred structure in subsequent layers. We presented models fit on both the CDR3 loop of α- and β-chain of the TCR (Fig. 1b) and models fit on the CDR3 loop of the β-chain and the antigen sequence (Fig. 2a). In both cases, we needed to integrate two sequences. To this end, we either used separate sequence-embedding layer stacks for each sequence (all models presented in Fig. 1 and models indicated as “separate” in Fig. 2) or by appending the two padded sequences and using a single sequence-embedding layer stack (models indicated as “concatenated” in Fig. 2). We reduced the positional encoding to a latent space of fixed dimensionality in the last sequence embedding layer of recurrent networks by the emitted state of the model on the last element of the sequence in each direction. This last layer allows usage of the same final dense layers independent of input sequence length. Convolutional and self-attention networks were not built to be independent of sequence length. We did, however, pad the input sequences to mitigate this problem on the data handled in this paper. We used a residual connection across all sequence-embedding layers. Further layer-specific hyper-parameters can be extracted from the code supplied in this manuscript (Supp. Data 1,2).

#### Final densely connected layers

We fed the activation generated in the sequence embedding block into a dense network that can integrate the sequence information with continuous or categorical donor- and cell-specific covariates. We modeled the binding event as a probability distribution over two states (bound and unbound) and compute the deviation of the model prediction from observed binding events via cross-entropy loss. Firstly, one can use such models to predict binding events on a single antigen represented as a single output node with a sigmoid activation function. Secondly, one can model a unique binding event among a panel of antigens with a vector of output nodes (one for each antigen and one node for non-binding) which are transformed with a softmax activation function.

#### Covariate processing

We set up a design matrix inspired by linear modelling to use as a covariate matrix. We modelled the donor as a categorical covariate, resulting in a one-hot encoding of the donor. We modelled total counts, negative control pMHC counts and surface protein counts as continuous covariates. We log(x+1) transformed negative control pMHC counts and surface protein counts to increase stability of training. We modelled total counts as the total count of mRNAs per cell divided by the mean total count.

### Train, validation and test splits

We used training data to compute parameter updates, validation data to control overfitting and test data to compare models across hyper-parameters. Model training was terminated once a maximum number of epochs was reached or if the validation loss was not decreasing any more. In the latter case, the model with the lowest validation in a sliding window of *n* epochs until the last epoch was chosen, *n* is given in the grid search scripts (Supp. Data 3). The model metrics presented in this manuscript are metrics evaluated on the test data. We provide training curves for all models that contributed to panels in this manuscript in Supp. Data 3.

### Optimization

We used the ADAM optimizer throughout the manuscript for all models. We used learning rate schedules that reduce the learning rate at training time once plateaus in the validation metric are reached. The initial learning rate and all remaining hyperparameters (batch size, number of epochs, patience, steps per epoch) were varied as indicated in the grid search hyperparameter list.

### Model fitting objectives

We chose cross-entropy loss on sigmoid or softmax transformed output activation values to train models that predict binarized binding events and mean squared logarithmic error (msle) on exponentiated output activation values for models that predict continuous (count) binding affinities.

### 10× CD8^+^ T-cell data processing

#### Primary data processing

We downloaded the full data of all four donors from^8^. All data processing for each model fit is documented in the package code (Supp. Data 1) and grid search scripts (Supp. Data 2). The number of T-cell clonotypes per antigen varied drastically between the order of 10^0^ and 10^4^ (Supp. Fig. 2a,b). Subsequently, we selected the 8 most common antigens (ELAGIGILTV, GILGFVFTL, GLCTLVAML, KLGGALQAK, RLRAEAQVK, IVTDFSVIK, AVFDRKSDAK, RAKFKQLL) for categorical panel model fits to avoid issues with class imbalances. We used the binarized binding event prediction by the authors of the data set^8^ (labeled “*_binder” in the files “*_binarized_matrix.csv”) as a label for prediction. For the continuous case, in which we predicted pMHC counts, we chose the corresponding count data columns in the same file. Next, we performed multiple layers of observations filtering: (1) doublet removal, (2) clonotype downsampling, and (3) class downsampling. It has previously been shown that doublets, i.e. droplets containing two cells targeted with the same barcode which cannot be distinguished in downstream analysis steps, tend to be enriched in subsets of transcriptome derived clusters^22^. We propose to use reconstructed TCR to identify potential doubles and demonstrate that the so characterized doubles are indeed enriched in a particular cluster in each donor (Supp. Fig. 1a-d). We further investigated the overall contribution of potentially ambient molecules that give rise to all observed T cells and found that high-frequency chains do not dominate the overall signal (Supp. Fig. 1e,f). This analysis presents an upper bound to the impact of ambient molecules on this experiment as evolutionary effects likely also contribute to over-representation of particular chain sequences. Subsequently, we removed all cellular barcodes that contain more than one α- or β-chain as mature CD8^+^ T cells are expected to only have a single functional α- and β-chain allele. Next, we down-sampled each clonotype to a maximum of 10 observations to avoid biasing the training or test data to large clones. Here, we used clonotypes as defined by the authors of the data set in the files “*_clonotypes.csv”^8^. Lastly, we downsampled the larger class to a maximum of twice the size of the smaller class when predicting a binary binding event for a single antigen. We did not perform this last step on multiclass and count prediction scenarios. We padded each CDR3 sequence to a length of 40 amino acids and concatenated these padded chain observations to a sequence of length 80 for models that were trained on both chains. We performed leave-one-donor-out cross-validation on models that did not take the donor identity as a covariate. We sampled 25% of the full data clonotypes and assigned all of the corresponding cells to the test set for all models that did use the donor covariate. The latter case yielded 68,716 clonotypes and 91,495 cells across all four donors. All cross-validations shown across different models are based on a 3-fold cross validation with seeded test-train splits resulting in the same split across all hyper-parameters.

#### Binarization of 10x CD8^+^ T-cell pMHC counts into bound and unbound states

We used the binarization described in the original publication^8^ for the raw counts to receive binary outcome labels: A total pMHC UMI count larger than 10 and at least five times as high as the highest observed UMI count across all negative control pMHCs was required for a binding event. If more than one pMHC passed these criteria, the pMHC with the largest UMI count was chosen as the single binder.

#### Test set assembly for models fit on IEDB data

This section describes how the test described in Fig. 2e and Supp. Fig. 5c was prepared. The cells were filtered as described above. We then extracted one binding TCR-antigen pair per cell from this list. We used the remaining TCR-antigen pairs as validated negative examples and down-sampled these to the number of positive observations to maintain class balance. All cross-validations shown across different models are based on a 3-fold cross validation with seeded test-train splits resulting in the same split across all hyper-parameters.

### IEDB data processing

#### Primary processing

We downloaded the data from the IEDB website^6^ with the following filters: linear epitope, MHC restriction to HLA-A*02:01 and organism as human and only human. This yielded a list of matched TCR (mostly β-chain CDR3s) with bound antigens. We assigned TCR sequences to a single clonotype if they were perfectly matched and downsampled all clonotypes to a single observation. We only extracted the β-chain and CDR3 sequences to a length of 40 amino acids. We padded the antigen sequences to a length of 25 amino acids. We sampled 10% of all observations as a test set. We generated negative samples for both training and test set separately by generating unobserved pairs of TCR and antigens. Here, we assumed that all TCRs bind a unique antigen out of the set of all antigen present in the database so that any other pairing would not result in a binding event. This procedure yielded 9,697 observations for both the positive and the negative set before the train-test split.

#### Test set assembly for models fit on IEDB data

This section describes how the test described in Supp. Fig. 5a was prepared. To explore the ability of antigen-embedding TcellMatch models to generalize to unseen antigens, we fit such a model on the subset of high-frequency antigens of IEDB with at least 5 unique TCR sequences and tested the models on the remaining antigens. All cross-validations shown across different models are based on a 3-fold cross validation with seeded test-train splits resulting in the same split across all hyper-parameters.

### VDJdb data processing

#### Primary processing

We provided an exploratory analysis of this data set in Supp. Data 3 “exploration_vdjdb_data.*”. We downloaded the data from the VDJdb^7^ website with the following filters: Species: human, Gene (chain): TRB, MHC First chain allele(s): HLA-A*02:Θ1. This yielded 3964 records. We assigned TCR sequences to a single clonotype if they were perfectly matched and downsampled all clonotypes to a single observation. We only extracted the β-chain and CDR3 sequences to a length of 40 amino acids. We padded the antigen sequences to a length of 25 amino acids.

#### Test set assembly for models fit on IEDB data

This section describes how the test described in Fig. 2d and Supp. Fig. 5b was prepared. We sub-selected observations with matching or non-matching antigens with respect to the training set depending on the application (described in the figure caption or main text). All cross-validations shown across different models are based on a 3-fold cross validation with seeded test-train splits resulting in the same split across all hyper-parameters.

